# A native herbaceous community exerts a strong allelopathic effect on the woody range-expander *Betula fruticosa*

**DOI:** 10.1101/2023.03.02.530791

**Authors:** Lichao Wang, Ayub M. O. Oduor, Yanjie Liu

## Abstract

Biological invasions by range-expanding native and alien plant species often reduce native plant community diversity and productivity. Superior performance of some invasive plants over native plants is due to production of allelochemicals by invaders that suppress growth of native plants. Nevertheless, native plants can also produce allelopathic compounds, which may provide biotic resistance against invasive plant species, in accordance with the homeland security hypothesis. In support of the hypothesis, several previous studies found evidence for allelopathic effects of native plant species on alien plant species. However, as most of these studies tested allelopathic effects of single native plant species on invasive plant species, the contribution of allelopathy to the resistance of native plant communities to invasion has received considerably less attention. Here, we performed two competition experiments in a greenhouse to test for potential pairwise allelopathic effects on each other of a woody range-expander *Betula fruticosa* and a community of four native herbaceous species in China. We tested whether *B. fruticosa* and the herbaceous community differed in their competitive effects and responses, and whether these were changed by the presence of activated carbon – an allelopathy neutralizer in the soil. Results show that presence of activated carbon ameliorated suppressive effects of the resident herbaceous community on above-ground biomass of *B. fruticosa*. By contrast, presence of activated carbon tended to aggravate suppressive effects of *B. fruticosa* on the resident herbaceous community. Overall, these results provide support to the homeland security hypothesis and suggest that strong biotic resistance of the resident herbaceous community may limit invasion success of the woody range-expander *B. fruticosa*.

## Introduction

Biological invasions by range-expanding native and alien plant species often have detrimental ecological and socioeconomic impacts (Essl et al. 2019; Schaffner et al. 2020; Vilà and Hulme 2017). As the number of range-expanding species will likely continue to increase (Essl et al. 2019; Seebens et al. 2021), it is important to identify mechanisms that underlie establishment of plants and invasibility of the resident plant communities. Invasive plants may outperform resident plants through superior competition for resources necessary for growth such as soil moisture, nutrients, and light (Gallien and Carboni 2017; Theoharides and Dukes 2007). Studies also indicate that superior performance of some invasive plants over native plants is due to production of allelochemicals by invaders that suppress growth of native plants (Baker 1974; Callaway and Aschehoug 2000; Hickman et al. 2021; Oduor et al. 2020). In support of this idea, a recent synthesis found that majority of 524 invasive plant species that were studied produce allelochemicals with the potential to negatively affect native plant growth (Kalisz et al. 2021). The observation that many invasive plants are allelopathic led to the formulation of the novel weapons hypothesis, which posits that some alien plants are successful invaders because they produce chemical compounds that are toxic to naïve native plants (Callaway and Ridenour 2004). Numerous studies find support for the novel weapons hypothesis (e.g., Abhilasha et al. 2008; Becerra et al. 2018; Inderjit et al. 2011; Ridenour and Callaway 2001; Thorpe et al. 2009), which suggests that allelopathy might indeed play an important role in plant invasions.

Native plants can also produce allelopathic compounds, which may provide biotic resistance against invasive plant species that are naïve to allelochemicals of the natives (Rabotnov 1982; Weidenhamer and Romeo 2005), in accordance with the homeland security hypothesis (Cummings et al. 2012). Several previous studies found evidence for allelopathic effects of native plant species on alien species (Adomako et al. 2019; Christina et al. 2015; Cummings et al. 2012; Hou et al. 2011; Mignoni et al. 2018; Ning et al. 2016; Weidenhamer and Romeo 2005; Yuan et al. 2021). However, as most of these studies tested allelopathic effects of single native plant species on invasive plant species, the contribution of allelopathy to the resistance of native plant communities to invasion has received considerably less attention (Yuan et al. 2022). Native plant communities may have strong allelopathic effects on invaders because there may be a high likelihood that species-rich native communities harbour at least one species that produces a disproportionately higher concentration of potent allelochemicals or there could be synergistic effects of different allelochemicals from different species within the community (Yuan et al. 2022). This prediction has received little empirical test (but see Adomako et al. 2019; Ning et al. 2016; Yuan et al. 2022).

Temperate herbaceous wetlands of China have been experiencing encroachment by various range-expanding shrub species in recent decades, as a consequence of global warming and anthropogenic disturbances (Lee et al. 2017; Lett and Dorrepaal 2018; Vuorinen et al. 2017). Because of the fundamental differences in functional traits between the range-expanding shrubs and native herbaceous species (Zhang et al. 2021a), shrub encroachment may threaten native plant community diversities. *Betula fruticosa* is one of the prominent range-expanding shrub species within the herbaceous wetlands (Zhang et al. 2021a). The species is distributed in Inner Mongolia and northern Heilongjiang (http://www.iplant.cn) where it grows at 600-1100 meters above sea level, and flowers and fruits between June and August. It has been expanding and gradually occupying a dominant position in the herb-dominated wetland habitats of the Sanjiang Plain in Northeast China (Zhang et al. 2021a). However, whether allelopathy mediates interaction between *B. fruticosa* and resident herbaceous communities remains unclear.

In this study, we performed a greenhouse experiment to test a prediction of the homeland security hypothesis that native plant communities may resist invasion by the range-expander *B. fruticosa* through inhibitory allelopathic effects. We addressed two questions: (1) Do *B. fruticosa* and the community have mutual allelopathic effects on each other? (2) Are the allelopathic effects of the community on *B. fruticosa* stronger than those of *B. fruticosa* on the community?

## Material and Methods

### Study species and cultivation

All the study species used for the experiment co-occur naturally in the Sanjiang Plain of northeast China, and their seeds were collected from wild populations (**Table 1**). On 3 January 2021, we sowed seeds of *B. fruticosa* into plastic trays (19.5 cm × 14.6 cm × 6.5 cm) that had been filled with a sterilized growth medium that comprised a mixture of sand and fine vermiculite (Pindstrup Plus, Pindstrup Mosebrug A/S, Denmark; pH: 6; 120.0 mg/L N; 12.0 mg/L P; 400.0 mg/L K; 28.0 mg/L Mg; 0.4 mg/L B; 2.0 mg/L Mo; 1.7 mg/L Cu; 2.9 mg/L Mn; 0.9 mg/L Zn; 8.4 mg/L Fe) 25%, 25%, 50%) in a ratio of 1:1 (v/v). The growth medium had been sterilized with a dose of 25 kGy of 60COγ irradiation for four days (at the Harbin Guangya Radiation New Technology Co., Ltd.,Harbin, China) to eliminate the potential influence of live soil organisms. One month later (i.e. 3 February 2021), we transplanted 200 similar-sized *B. fruticosa* individuals into a seedling tray, one individual in each tray hole. Between 19 February and 23 March, we fertilized the seedlings with 1g/L of Peters Professional® liquid fertilizer once every week for a total of five weeks. The fertilizer contained the following nutrients: 20% nitrogen, 20% potassium oxide (K_2_O), 0.02% Boron, 0.015% Cooper, 0.12% Iron, 0.06% Manganese, 0.010% Molybdenum, and 0.08% Zinc. On 15 February, we sowed approximately 200 seeds for each of the four wetland herb species in trays that had been filled with the same sterilized growth medium as above. We placed all trays with seeds in a greenhouse under natural light conditions, with a temperature between 15 and 25 ℃.

**Table 1.**
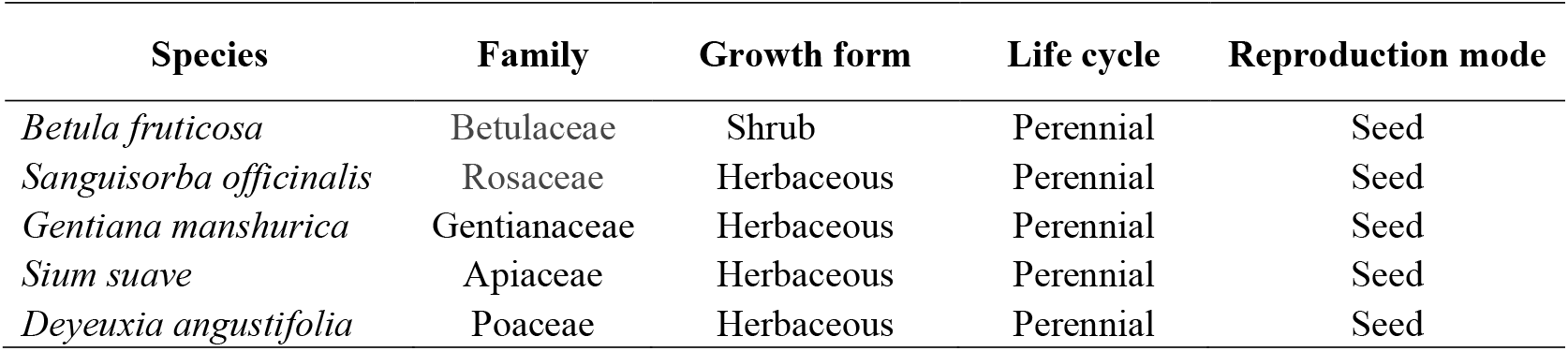
Information on the plant species that were used in the present study

### Experimental set-up and measurement

To assess whether allelopathy mediates competitive interactions between *B. fruticosa* and the resident native herbaceous community, we performed two separate greenhouse experiments. The first experiment aimed to test allelopathic effect of the woody plant *B. fruticosa* on the herbaceous community of native species (Experiment-1), while the second experiment tested allelopathic effect of the herbaceous community on *B. fruticosa* (Experiment-2) (**Figure S1**). Both experiments employed a fully-crossed factorial design. In Experiment 1, we grew the native community with vs. without *B. fruticosa* and in the presence vs. absence of activated carbon (**Figure S1**). In Experiment 2, we grew the woody plant *B. fruticosa* with vs. without the native community and in the presence vs. absence of activated carbon (**Figure S1**). Activated carbon is widely used in ecological studies to adsorb and neutralize potential allelochemicals produced by experimental plants (Kabouw et al. 2010; Zhang et al. 2021b). To set up the experiments, we filled 160 circular 2.5-L plastic pots (top diameter × bottom diameter × height: 18.5 cm × 12.5 cm × 15 cm) with a sterilized growth medium that had been prepared as described above. In each pot, we homogenized the growth medium with 5 g of a slow-release fertilizer (Osmocote Exact Standard; 15.00% N + 9.00% phosphorus pentoxide + 12.00% potassium oxide + 2.00% magnesium oxide + 0.02% B + 0.05% Cu + 0.45% Fe + 0.09% cheated by EDTA + 0.06% Mn + 0.02% Mo + 0.015% Zn; Everris International B.V., Geldermalsen, The Netherlands). We then homogenized pot content with 50 mL of activated carbon (Analytical purity of activated carbon powder, pH 5.0-7.0; Sinopharm Chemical Reagent Co., Ltd, China), for a half of the pots (n=80).

In Experiment 1, we transplanted an individual of each of the four native herbaceous species in 80 pots. A half of the pots contained activated carbon, while the other half did not. The individual seedling were transplanted at equal distances from each other in a circular formation. Then we introduced an individual *B. fruticosa* into the middle of the pot, for a half of the pots with and without activated carbon. In experiment 2, we transplanted *B. fruticosa* individuals in the center of the pot. In a half of pots with and with activated carbon treatment, we transplanted an individual seedling of each of the four herbaceous species at equal distances from each other in a circular formation around *B. fruticosa*. Each of the two experiments had two levels of competition: (Community grown without *B. fruticosa* vs. community grown with *B. fruticosa* for Experiment 1 and *B. fruticosa* grown with community vs. *B. fruticosa* grown without community for Experiment 2) × 2 levels of activated carbon (with vs. without activated carbon) × 20 replicates (**Figure S1**). Immediately after transplant, we randomly assigned the 160 pots to positions on one greenhouse bench (temperature: 22-28 ℃; relative humidity 60%; natural lighting) and watered them regularly. The two experiments lasted 123 days and were performed concurrently at the Northeast Institute of Geography and Agroecology of the Chinese Academy of Sciences (43°59’49”N, 125°24’3”E).

On 2 July 2021, we harvested above-ground biomass of native herbaceous community in Experiment 1 and above-ground biomass of *B. fruticosa* in Experiment 2. Three *B. fruticosa* individuals (one grown alone) died during the experiment, thus we harvested above-ground biomass of 77 *B. fruticosa* and 80 herbaceous community. All the plant materials were dried at 65 ℃ for 72 hours and then weighed to an accuracy of 0.0001 g.

### Statistical analyses

To test the mutual allelopathy between *B. fruticosa* and the native herbaceous community, we fitted two separate Bayesian multilevel models in R 4.0.2 (R Core Team 2020) using the *brm* function in the *brms* package (Bürkner 2017). As the above-ground biomass of *B. fruticosa* and the native herbaceous community had Gaussian error distributions, we applied cube root transformation of the data prior to the analysis to improve normality and homogeneity of the residuals. The fixed part of each model included competition treatment (two levels each for experiments 1 and 2), activated carbon treatment (with vs. without activated carbon), and two-way interaction between the two factors. Each model was run with four independent Markov chains of 4, 000 iterations, discarding the first 2, 000 iterations per chain as warm-up and resulting in 8, 000 posterior samples overall. To directly test hypotheses about the main and interactive effects based on each coefficient’s posterior distribution, we used the sum coding, which effectively ‘centers’ the effects to the grand mean (i.e., the mean value across all data observations; Schad et al. 2020). To implement this in *brms*, we used the functions *contrasts* and *contr*.*sum* of the *stats* package in R. We considered the two fixed factors (competition and activated carbon), and interaction between them as significant when their 95% credible interval of the posterior distribution did not overlap zero, and as marginally significant when their 90% credible intervals did not overlap zero.

## Results

Above-ground biomass of *B. fruticosa* was significantly influenced by main and interactive effects of competition and activated carbon treatments (**Table 2; Fig. 1**). In the absence of activated carbon, competition from the community reduced above-ground biomass of *B. fruticosa* by 68.43% (**Fig. 1a**). However, in the presence of activated carbon, *B. fruticosa* produced similar above-ground biomass in the presence (1.16 g) vs. absence (1.32 g) of competition from the herbaceous community (**Fig. 1a**). By contrast, above-ground biomass of the resident herbaceous community was significantly influenced by the main effect of competition from *B. fruticosa* and marginally by an interaction between activated carbon and competition (**Table 2; Fig. 1)**. Specifically, averaged across the activated carbon treatments, competition from *B. fruticosa* reduced mean above-ground biomass of the herbaceous community by 19.4% (**Fig. 1b**). Nevertheless, presence of activated carbon marginally aggravated the suppressive effect of *B. fruticosa* on the community above-ground biomass (**Fig. 1c**). Specifically, competition from *B. fruticosa* reduced the community above-ground biomass by 30.33% in the presence of activated carbon and by 5.7% in the absence of activated carbon (**Fig. 1c)**.

**Table 2.** Output of two separate Bayesian multilevel models that were run to test the effects of competition treatment (Community grown without *B. fruticosa* vs. community grown with *B. fruticosa* for Experiment 1 and *B. fruticosa* grown with community vs. *B. fruticosa* grown without community for Experiment 2), activated carbon treatment (with vs. without activated carbon), and two-way interaction between the two treatments on above-ground biomass of *B. fruticosa* and a resident herbaceous community in each pot. Shown are the model estimates and standard errors (SE) as well as the lower (L) and upper (U) values of the 95% and 90% credible intervals (CI).

|  |  | Estimate | SE | L95%CL | U95%CL | L90%CL | U90%CL |
| --- | --- | --- | --- | --- | --- | --- | --- |
| **Above-ground biomass of *B. fruticosa*** | Intercept | 0.958* | 0.053 | 0.854 | 1.061 | 0.869 | 1.045 |
|  | Activated carbon (A) | 0.114* | 0.053 | 0.01 | 0.218 | 0.027 | 0.202 |
|  | Competition (C) | 0.130* | 0.052 | 0.027 | 0.233 | 0.044 | 0.217 |
|  | A×C | 0.128* | 0.054 | 0.021 | 0.236 | 0.04 | 0.217 |
| **Above-ground biomass of resident herbaceous community** | Intercept | 1.796* | 0.025 | 1.748 | 1.844 | 1.755 | 1.837 |
|  | Activated carbon (A) | -0.008 | 0.024 | -0.056 | 0.04 | -0.048 | 0.033 |
|  | Competition (C) | 0.071* | 0.025 | 0.022 | 0.119 | 0.029 | 0.111 |
|  | A×C | -0.048 † | 0.024 | -0.095 | 0.001 | -0.087 | -0.008 |

Model estimates whose 95% credible intervals do not overlap with zero are indicated with asterisks (\*), and those whose 90% credible intervals do not overlap with zero are indicated with daggers (†).

**Figure 1.**
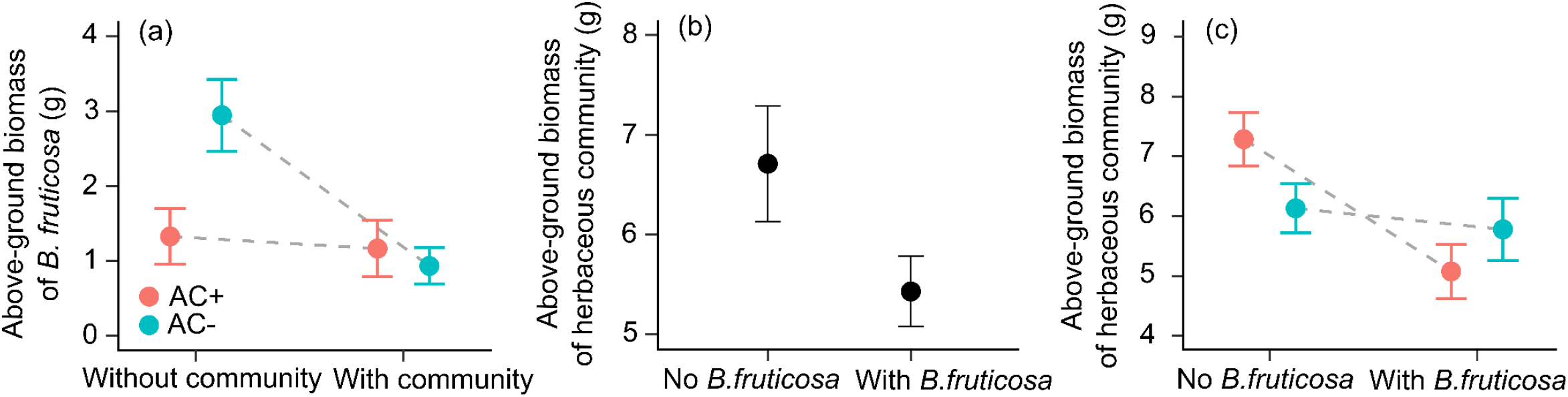
Mean values (± SE) of above-ground biomass of *B. fruticosa* (**a**), herbaceous community (**b & c**). Shown are main and interactive effects of presence (with community) vs. absence (without community) of competition from a resident herbaceous community, presence (AC+) vs. absence (AC-) of activated carbon, and presence (with *B. fruticosa*) vs. absence (No *B. fruticosa*) of competition from *B. fruticosa*(*cf*. **Table 2**).

## Discussion

We tested allelopathic interactions between a range-expanding native shrub *B. fruticosa* and a native herbaceous community, and results suggest that presence of activated carbon ameliorated suppressive effects of the resident herbaceous community on above-ground biomass of *B. fruticosa* (**Fig. 1a**). By contrast, presence of activated carbon tended to aggravate suppressive effects of the range-expander shrub on the resident herbaceous community (**Fig. 1c**). Overall, these results suggest that the resident herbaceous community had a strong allelopathic effect on *B. fruticosa*, and that these effects were neutralized or reduced by activated carbon. On the other hand, *B. fruticosa* had little allelopathic effect on the community. Taken together, the results provide a strong support for the homeland security hypothesis (Cummings et al. 2012), but little support for the novel weapons hypothesis (Callaway and Ridenour 2004).

The present results complement those of the few other studies that tested whether the novel-weapons hypothesis and homeland-security hypothesis can both explain competitive interactions between alien and native plant species within the same community, with mixed findings. While the novel weapons hypothesis predicts strong allelopathic effects of alien plant species on native plant species (Callaway and Ridenour 2004), the homeland security hypothesis posits that the reverse should also be true (Cummings et al. 2012). A recent study of interactions between five alien and five native herbaceous species in China found that alien species had negative allelopathic and resource-based competitive effects on native plants, while the reverse was also true (Yuan et al. 2021). The native plant *Sesbania virgata* had strong allelopathic effects on the alien plant *Leucaena leucocepahala* in Brazil, while the alien did not affect the native through allelopathy (Mignoni et al. 2018). Other studies have demonstrated allelopathic effects of native plant species on alien species but did not simultaneously test allelopathic effects of aliens on natives (e.g., Cummings et al. 2012; Hou et al. 2011; Weidenhamer and Romeo 2005). More studies are needed to enhance understanding of the generality of both novel weapons hypothesis and homeland security hypothesis operating in the same invaded communities.

Our result showed that suppressive effects of *B. fruticosa* on the native herbaceous community tended to be stronger in the presence (rather than absence) of activated carbon (**Fig. 1c**). This could partly reflect that activated carbon released the allelopathic suppression among native species in the non-invaded community, and thus increased their biomass production. On the other hand, activated carbon tended to facilitate greater growth of *B. fruticosa* in the invaded community, and hence it’s stronger suppressive effects on the community. Activated carbon has been successfully used in several studies on allelopathy (e.g., Inderjit and Callaway 2003; Lankau 2010; Mahall and Callaway 1992; Mangla, Callaway 2008; Yuan et al. 2021). However, activated carbon can alter soil nutrient availability and plant growth even in the absence of the focal allelopathic agent (e.g., Kabouw et al. 2010; Lau et al. 2008; Weißhuhn and Prati 2009). For example, activated carbon caused an increase in the specific root length and root branching in a study of 10 plant species (Yuan et al. 2021). In the present case, it is likely that activated carbon caused a reduction in soil nutrient availability to plants by adsorbing the nutrients leading to stronger competitive effects of *B. fruticosa* on the resident herbaceous community.

The present results suggesting significant negative allelopathic effects of a native plant community on the range-expander *B. fruticosa*, complement those of a few other studies that tested for allelopathic effects of native plant communities on alien plant species, with mixed results. For instance, native plant communities had negative allelopathic effects on the invader *Solidago canadensis* in China (Adomako et al. 2019). Ning et al. (2016) also found evidence for allelopathic effects of native plant communities on alien species in Germany. These findings provide support for the homeland-security hypothesis. In contrast, allelochemicals of diverse native plant communities did not provide resistance but instead facilitated the germination of six alien invasive plant species (Yuan et al. 2022). However, as the few studies that have tested allelopathic effects of native plants on invasive plants have focused on herbaceous invasive species only, it remains unclear whether allelopathic effects of native plant communities may differ between woody and non-woody invasive species.

In conclusion, our results suggest an asymmetric allelopathic interaction between the native herbaceous community and *B. fruticosa* as the native herbaceous community likely had strong negative effect on *B. fruticosa* through alleloptahy, but the reverse was not true. In line with the homeland security hypothesis, strong biotic resistance of the resident herbaceous community may limit invasion success of the range-expander *B. fruticosa*.

## Acknowledgements

We thank Xue Zhang, Huifei Jin and Mingxin Pan for help with the set-up of the experiment and plant harvest.

## Author contributions

YL conceived the idea and designed the experiment. LW performed the experiment. LW and YL analyzed the data. LW wrote the draft of the manuscript, with further inputs from YL and AMOO.

## Data accessibility

Should the manuscript be accepted, the data supporting the results will be archived in Dryad and the data DOI will be included at the end of the article.

## Supporting information

**Figure S1.**
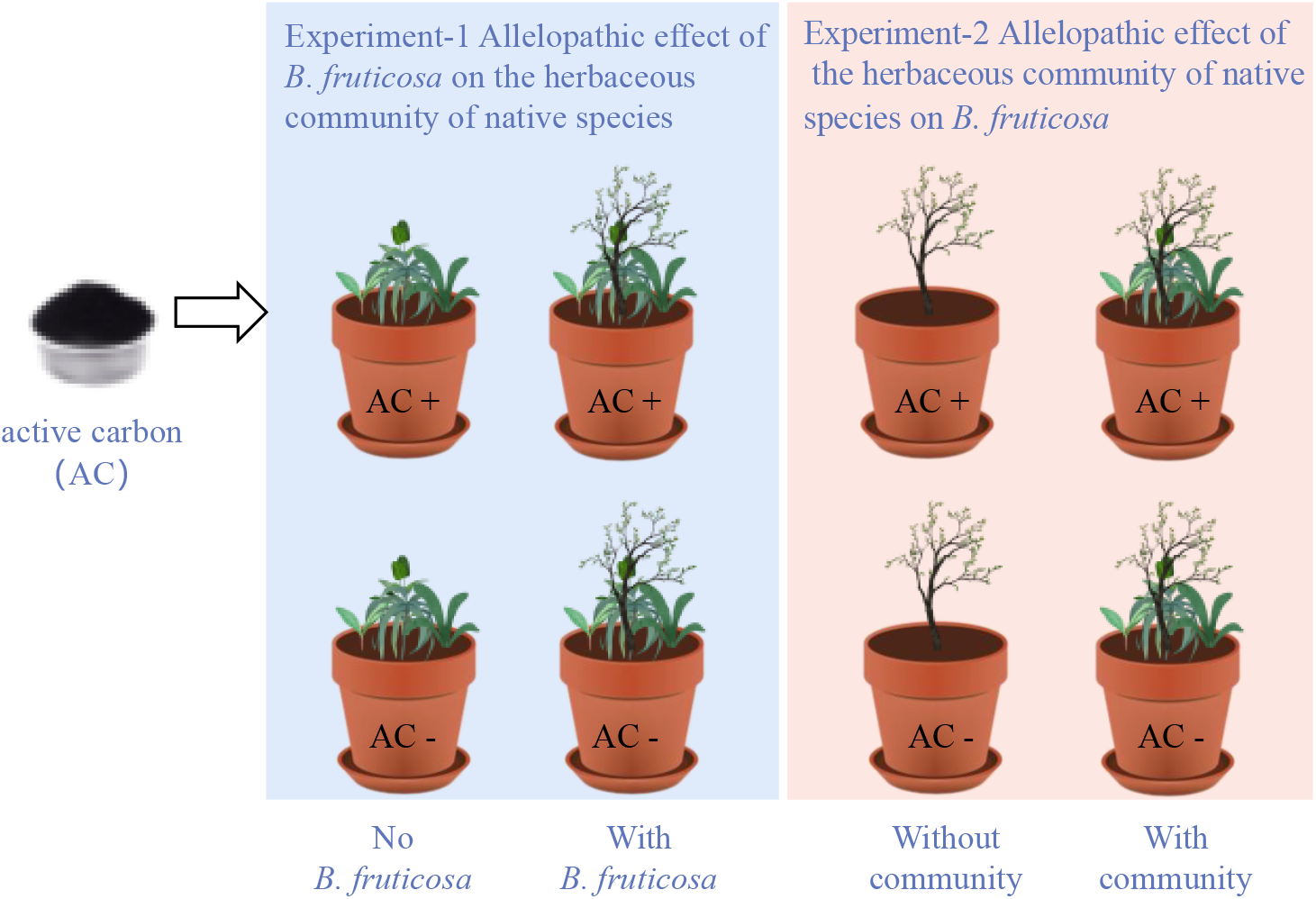
A schematic of an experimental set-up to test allelopathic interactions between a range-expanding species *Betulla fruticosa* and a native plant community of four species *Sanguisorba officinalis, Gentiana manshurica, Sium suave*, and *Deyeuxia angustifolia*. Activated carbon was used to neutralize allelochemicals.

